# Distinct associative learning abilities for colour and odour in the flower-feeding *Drosophila elegans* and the fruit-feeding *Drosophila melanogaster*

**DOI:** 10.64898/2026.09.24.753984

**Authors:** Kohei Kawamura, Christian König, Teiichi Tanimura, Bertram Gerber, Yuki Ishikawa

## Abstract

Animal behaviour is both innately constrained and shaped by learning. This mosaic organization has evolved in response to species-specific ecological demands and may differ between sensory modalities. Flower-visiting animals are a particularly useful system for investigating the relationship between sensory ecology and learning because they rely on multiple floral cues, particularly odour and colour, to locate food sources. However, it remains largely unexplored whether specialization on floral resources entails divergence in learning abilities across sensory modalities. *Drosophila elegans* is a flower-feeding species that depends heavily on floral resources throughout its life; adults spend much of their time on flowers and larvae develop on fallen flower leaves. Here, we compared odour-reward and colour-reward associative learning between the flower-feeding *D. elegans* and the fruit-feeding *D. melanogaster*. We found that, under conditions of equilibrated motivation, odour- and colour-preference, and using the same sugar reward, *D. elegans* exhibited poorer odour-reward learning performance but better colour-reward learning performance than *D. melanogaster*. These results suggest that the modality-specific eligibility of sensory information to enter into associations, known as the ‘Garcia-effect’ in experimental psychology, can evolve oppositely between species. This highlights the relationship between ecological specialization and mnemonic processing, and shows that biological ‘intelligence’ is not general.

## 1. Introduction

Animal foraging is shaped by both innate and learned behavioural tendencies. While innate preferences can predispose animals to respond to specific food-related cues, associative learning enables them to link predictive stimuli with resource availability based on their individual experience and to adjust individual food search according to local ecological conditions ^1–3^. Because different species face different challenges in locating, acquiring, and memorizing food sources, their associative learning abilities are expected to have evolved to meet these species-specific ecological demands. Several studies have shown that closely related species differ in learning abilities relevant to foraging and/or feeding. For instance, insect- and fruit-feeding bats exhibit superior taste-learning abilities compared with blood-feeding species ^4^, and diurnal frogs possess better spatial learning abilities than nocturnal species ^5^. These examples suggest that associative learning abilities can evolve according to ecological demands to improve foraging efficiency and enhance fitness.

Feeding on floral resources, such as nectar, pollen, or the flower leaves themselves, has evolved across diverse animal taxa and is common among insects ^6–8^. Many flower-visiting insects act as pollinators, mediating key plant-animal interactions in terrestrial ecosystems ^9,10^. Indeed, flowers ‘actively’ attract insects by providing distinctive visual and olfactory signals indicative of the resources they offer ^11–15^. To exploit these signals, flower visitors use sensory systems adapted to detect, process and memorize these signals^13,16–18^.

The quality and quantity of floral resources may vary among plant species, individual plants, habitat and time of day, and thus flower-visiting animals can improve their foraging efficiency by learning the association between floral traits and the reward available from flowers ^1,19–23^. Fittingly, associative learning of floral signals in bumblebees and hawk moths has been shown to facilitate visits to flowers that provide greater rewards ^21,22^. Honeybees can likewise integrate information from multiple sensory modalities during associative learning ^19,23^. However, whether the relative ease at which stimuli from different sensory modalities are learned as predictors, known from experimental psychology as the ‘Garcia effect’ ^24^, is differentially shaped by evolution in different species remains largely unexplored.

Associative learning has been extensively studied in *Drosophila melanogaster* ^25,26^. Robust behavioural assays using odours and colours as conditioned stimuli have been developed, revealing the neural and molecular mechanisms underlying associative learning. *D. elegans* is a closely related species that diverged from *D. melanogaster* approximately 15 million years ago, and represents a species group (*elegans* group) distinct from the *melanogaster* species group (Fig. 1) ^27,28^. Unlike the fruit-feeding *D. melanogaster*, *D. elegans* is strongly reliant on flowers throughout much of its life history^27^. Males establish territories and court and mate with females within flowers, females oviposit there, and larvae develop by feeding of fallen flowers ^29–31^. These observations suggest a strong dependence on floral resources, making *D. elegans* a useful system for testing whether associative learning abilities have evolved with floral-resource use. In *D. melanogaster*, orientation toward food is known to depend on a combination of olfactory stimuli and achromatic visual contrasts ^32,33^, and in otherwise similar paradigms associative learning performance is characteristically better for olfactory than for visual stimuli ^26,34^. In *D. elegans*, little is known about the sensory cues used for locating flowers, or how easily stimuli from different sensory modalities can enter into association with reward. What is known, however, is that this species visits a variety of flowers with diverse visual and olfactory characteristics ^27,29,35^. Accordingly, the sensory information relevant for resource localization and memory formation in *D. elegans* may differ from the information that is relevant to *D. melanogaster*.

**Fig. 1.**
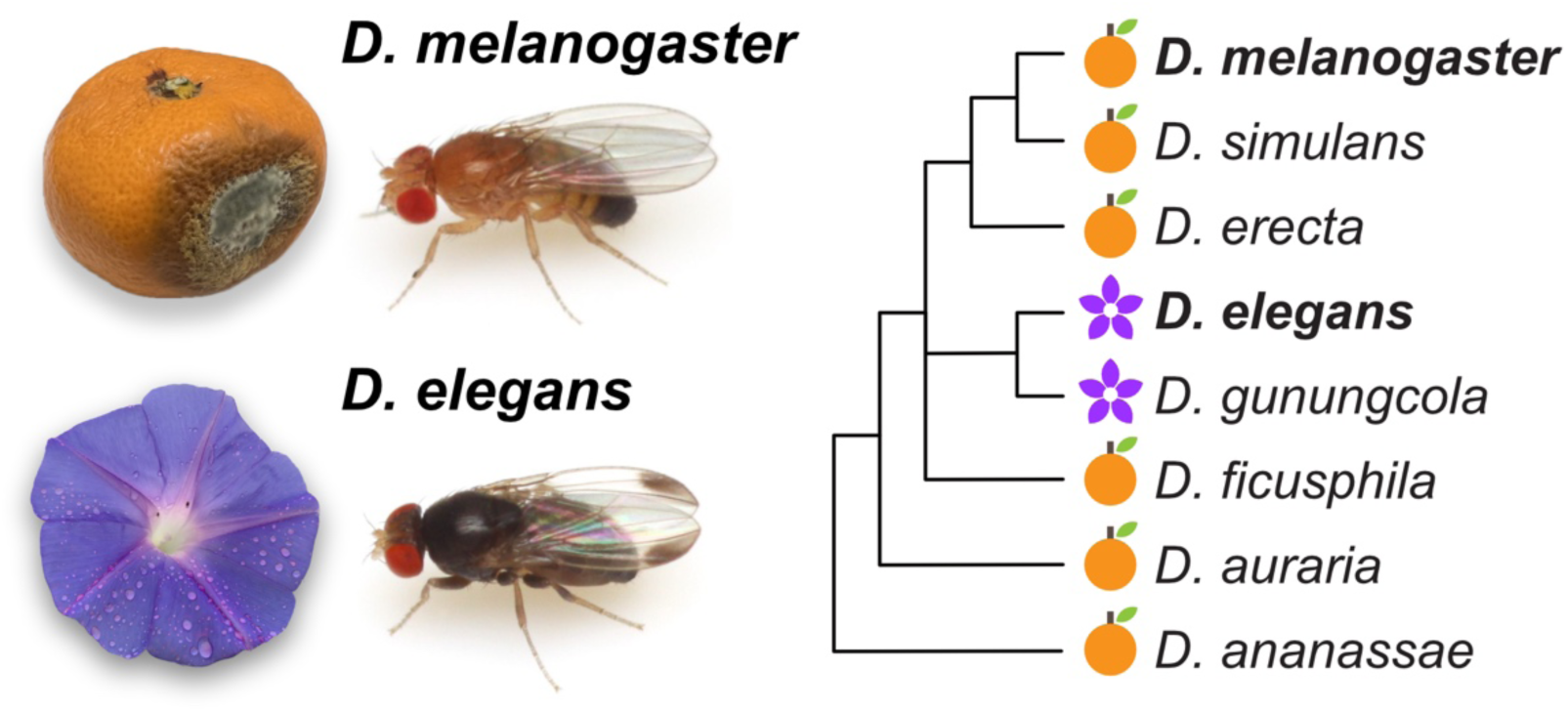
Fruit-feeding *Drosophila melanogaster* (left, top) and flower-feeding *D. elegans* (left, bottom) (both male). A mandarin orange and a morning glory flower, shown not to scale, serve as examples of their respective food resources. The phylogenetic relationships among species in the genus *Drosophila* are shown to the right (modified from ^27^).

Here, we compared odour- and colour-reward associative learning between the flower-feeding species *D. elegans* and the fruit-feeding species *D. melanogaster*. If specialization on floral resources has evolutionarily influenced learning abilities, we predicted that associative learning performance would differ between these two species. In particular, we asked whether between-species differences in learning performance are consistent across odour- and colour-learning.

## 2. Material and Methods

### (a) Stocks and fly maintenance

We used 3–8-day-old male and female adult flies of the wild-type *D. melanogaster* strain *Canton-S*, and the wild-type *D. elegans* strain IR-MTK (originally collected on Iriomote Island, Okinawa, Japan, and distributed by Ehime-Fly stock centre). All flies were maintained at 25 ℃ and approximately 65% relative humidity under a 12 h light/12 h dark cycle. Flies were reared on fly food containing 3.15% yeast, 2% raisins, 0.125% honey, 0.125% cane syrup, 2.5% semolina, 0.417% agar (w/v) and were randomly selected for experiments as mixed-sexed cohorts. Fresh food was provided every 2-3 days.

### (b) Starvation resistance

Newly eclosed flies were maintained on vials with fly food. Groups of 15-25 individuals aged 3 days were then transferred to new vials containing moist tissue paper and maintained at 25°C and 65% relative humidity (N = 10 vials for each species). The number of dead flies was recorded twice each day, at 9:00 and 16:00, until all individuals had died.

### (c) Setup for behavioural experiments

Behavioural experiments used a T-maze setup from CON-ELEKTRONIK (Greussenheim, Germany) placed in an environmental cabinet maintained at 25°C and > 40% relative humidity ^36^. To control for potential positional effects within the setup, the assignment of stimuli to the two arms was counterbalanced whenever applicable.

### (d) Innate sucrose preference

Chromatography paper (3MM Chr, WHA3030917, Whatman) treated with either 2 M sucrose solution (S0389, Sigma-Aldrich) or with tap water and dried overnight (hereafter referred to as sucrose paper and water paper, respectively) was placed along the choice arms of the setup. Approximately 20-60 experimentally naïve flies starved for 0, 24 or 48 hours were introduced to the central compartment of the setup and shortly thereafter released into its choice arms. After 2 min, the numbers of flies present in the sucrose and water arms were recorded. A sucrose preference index was quantified as (*N_Sucrose_* – *N_Water_)* / (*N_Sucrose_* + *N_Water_*). The numbers of independent experiments were 4, 12, and 15 for *D. elegans*, and 4, 9, and 2 for *D. melanogaster* at 0, 24, and 48 hours of starvation. The smaller number of experiments at 48 hours in *D. melanogaster* was due to the relatively early mortality upon starvation in this species.

### (e) Starvation treatment

Prior to the experiments measuring innate odour preference, innate colour preference, and prior to the learning experiments, flies were starved as described above for 24 and 72 hours in *D. melanogaster* and *D. elegans*, respectively. These durations were determined based on the starvation-resistance assay described above.

### (f) Innate odour preference

Innate odour preferences for 1-octanol (OCT) (8.20931, Sigma-Aldrich) *versus*benzaldehyde (BA) (8.01756, Sigma-Aldrich) was examined by applying 250 μl and 50 μl of these odorants, respectively, to the 1 cm-deep Teflon containers of the setup, at 14 mm and 5 mm diameter, respectively ^36^. Untreated chromatography paper was placed along the inner wall of each arm of the setup. OCT and BA were presented in the opposite arms. Approximately 30-60 experimentally naïve flies were introduced to the central compartment of the setup and shortly thereafter released into its choice arms. After 1 min, the numbers of flies in the OCT and BA arms were recorded. An odour preference index was calculated as (*N_OCT_* – *N_BA_)* / (*N_OCT_* + *N_BA_*). The numbers of independent experiments were 24 for *D. elegans*, and 19 for *D. melanogaster*.

### (g) Odour-reward associative learning

Flies were trained using OCT or BA as conditioned stimuli (CS) and sucrose as the unconditioned stimulus (US). Learning performance was quantified in a test phase in which flies were allowed to choose between the two odorants. All training sessions were initiated between ZT1 and ZT7. The numbers of independent experiments were 30 for *D. elegans*, and 31 for *D. melanogaster*.

#### Training phase

Approximately 30-60 experimentally naïve flies were introduced to the central compartment of the setup and shortly thereafter released into its training arm. Either OCT or BA was designated as the CS+, whereas the alternative odorant served as the CS−. Therefore, the assignment of OCT and BA as CS+ or CS− was counterbalanced across reciprocal training groups. Flies were first exposed for 3 min to the CS+ odorant presented in an arm lined with sucrose paper. They were then gently collected into empty vials using an aspirator; this procedure of collecting the flies was necessary because *D. elegans* tended to cling tightly to the filter paper and could not be reliably dislodged by the tapping procedure routinely used for *D. melanogaster* (in all present learning experiments, flies from both species were transferred with the aspirator). Following an 8 min interval, the flies were reintroduced into the setup and exposed for 3 min to the CS− odour presented in an arm lined with water paper. Thereafter, flies were collected as described.

#### Test phase

Following training, flies were allowed to choose between OCT and BA. Untreated chromatography paper was placed along the inner wall of each arm. OCT and BA were presented in opposite arms during the test. 10 min after training, flies were introduced to the central compartment of the setup and shortly thereafter released into its choice arms. After 1 min, the numbers of flies in the OCT and BA arms were recorded. Only trials with more than 10 flies remaining at the end of the experiments were included in subsequent analyses.

Odour preference indices (PIs) were calculated from the raw data, then learning indices (LIs) were calculated as (PI_CS+_ – PI_CS−_)/2 × 100 for each pair of reciprocally trained groups of flies. Positive LI values indicate a learned, associative preference for the previously rewarded odour.

### (h) Visual stimuli

Blue and green light stimuli were presented by LED devices designed to fit the setup (CON-ELEKTRONIK, Greussenheim, Germany). Each device consisted of a hollow cylinder with an outer diameter of approximately 50 mm and an inner diameter of approximately 30 mm. Blue or green LEDs were embedded in the inner wall of the cylinder in a 3 × 8 array. Light intensity was adjusted using a 16-bit pulse-width modulation controller that allows precise regulation of LED output.

To ensure that flies learned colour rather than brightness differences, the subjective brightness of the two colours, defined as the effective light intensity to the fly retina, was matched between blue and green light. First, the emission spectra of all LED devices used in the experiments were measured five times each using an STS-VIS miniature spectrometer (Fig. S1a; product no. S05690, Ocean Optics). Subjective brightness was calculated from the measured emission spectra and the spectral sensitivity of the fly retina measured by Katsura et al. (unpublished data). The intensity of each LED was adjusted for each species such that blue and green light produced indistinguishable subjective brightness (Fig. S1b). No significant difference in subjective brightness was detected between blue and green light for either species (one-way ANOVA followed by Tukey’s post hoc test, Table S1).

### (i) Innate coloured-light preference

Innate coloured-light preference for blue *versus* green light was examined. Coloured-light stimuli were presented by attaching the LED devices to the outside of the arms of the setup. Untreated chromatography paper was placed along the inner wall of each arm. 30-60 experimentally naïve flies aged 6-8 days were introduced to the central compartment of the setup and briefly thereafter released into its choice arms. After 1 min, the numbers of flies in the blue- and green-light arms were recorded. Colour preference indices were calculated as (*N_Blue_* – *N_Green_)* / (*N_Blue_* + *N_Green_*). The numbers of independent experiments were 26 for *D. elegans*, and 20 for *D. melanogaster*.

### (j) Colour-reward associative learning

Flies were trained using blue and green light as CS. Learning performance was quantified in a test phase in which flies were allowed to choose between the two colours. The numbers of independent experiments were 29 for *D. elegans*, and 27 for *D. melanogaster*.

#### Training phase

The experimental setup and procedures were otherwise identical to those used in the odour-reward associative learning assay, except for the modifications described below. Either blue or green light was designated as the CS+, whereas the alternative light served as the CS−. Flies were first exposed for 3 min to the CS+ light presented in an arm lined with sucrose paper. Following an 8 min interval, the flies were reintroduced to the setup and exposed for 3 min to the CS− light presented in an arm lined with water paper. As in the odour-reward associative learning assay, reciprocal training groups were used, with blue and green light alternately assigned as CS+ and CS−.

#### Test phase

Following training, flies were allowed to choose between blue and green light. Blue and green light were presented in opposite choice arms during the test. 10 min after training, flies were introduced to the central compartment of setup and shortly thereafter released into its choice arms. After 1 min, the numbers of flies in the blue and green arms were recorded. Colour preference indices and LIs were calculated, with due modifications, as described above.

### (k) Statistical analyses

All data processing, calculations, and statistical analyses were conducted in R version 4.5.1 (R Core Team, 2025).

For starvation resistance, survival curves were estimated using the Kaplan–Meier method, and median survival times (LT50) were calculated using the *survival* package (ver. 3.8.6). Differences in survival between species were assessed using log-rank tests. For sucrose preference, odour preference, odour-reward associative assay, coloured-light preference, and colour-reward associative assay, normality and homogeneity of variance were assessed using Shapiro–Wilk and Bartlett tests, respectively. Depending on whether the assumptions of parametric tests were met, data were analysed using Wilcoxon signed-rank tests, Wilcoxon rank-sum tests, Student’s t-tests, one-way analysis of variance (ANOVA), or aligned rank transform ANOVA (ART ANOVA) (see captions of each figure and table). Post hoc comparisons were performed using Tukey’s honestly significant difference test or estimated marginal means, as appropriate. P-values were adjusted for multiple comparisons using the Holm method when applicable. ART ANOVA and estimated marginal means were implemented using the R packages *ARTool* (ver. 0.11.2) and *emmeans* (ver. 2.0.1), respectively.

## 3. Results

To compare the learning abilities of *D. elegans* and *D. melanogaster*, we conducted an odour–reward associative learning assay using a T-maze paradigm ^36^. As this assay requires starvation prior to training ^37,38^, we determined the appropriate starvation durations in both species. In *D. melanogaster*, survival declined sharply approximately 30 hours after the onset of starvation, with a median survival time (LT50) of 46 hours (Fig. 2a). In contrast, *D. elegans* exhibited higher starvation resistance; survival remained high until approximately 80 hours after starvation began, and LT50 was 118 hours. Based on these results, we starved *D. melanogaster* for 24 hours and *D. elegans* for 72 hours in all subsequent experiments (unless noted otherwise) as these starvation durations correspond to the period immediately before survival began to decline in the respective species, thus equilibrating both species for motivational state.

**Fig. 2.**
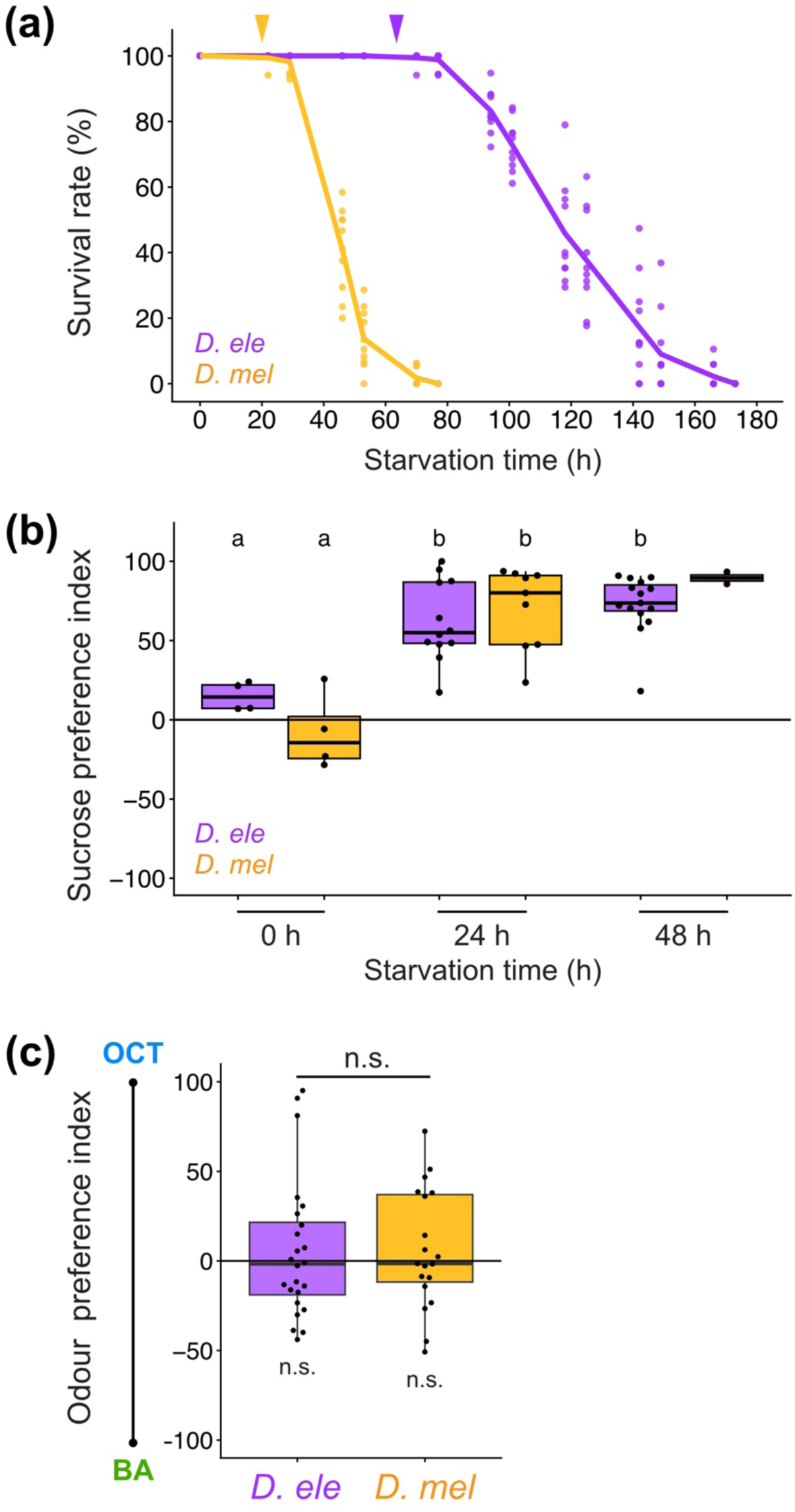
(a) *D*. *elegans* is more resistant to starvation than *D*. *melanogaster* (log-rank test, χ² = 1794, df = 1, p < 2 × 10⁻¹⁶, N = 10 replicates for each species). Dots represent the survival rate of flies in each vial at each time point. Lines connect the mean survival rates across vials. Arrowheads indicate the starvation durations used in the subsequent learning assays. (b) Starvation increased sucrose preference in both species, with no significant interspecific difference detected (see Table S2, N = 4, 12, and 15 for *D. elegans*, and 4, 9, and 2 for *D. melanogaster* at 0, 24, and 48 h of starvation, respectively). Different letters above each individual bar indicate statistically significant differences (ART ANOVA, p < 0.05). (c) Innate odour preference for octanol (OCT) *versus* benzaldehyde (BA). Neither species exhibited a significant preference for either odour (Wilcoxon signed-rank test: *D*. *elegans*, p = 1, N = 24; *D*. *melanogaster*, p = 0.822, N = 19; n. s. placed below individual bars). No significant difference was detected between species (Wilcoxon rank-sum test, p = 0.651; n.s. placed above bars).

Since responsiveness to sucrose, which served as the reward in the associative learning assay, can vary among genotypes and influence learning performance ^39,40^, interspecific differences in sucrose responsiveness could potentially confound comparisons of learning ability. We therefore compared innate sucrose preference between the two species under different starvation conditions. Starvation induced a significant preference for sucrose in both species, and sucrose preference did not differ between species across the starvation treatments examined (Fig. 2b, Table S2 and S3).

We further tested for possible interspecific differences in their innate odour preference between octanol *versus* benzaldehyde, the odours to-be used in the learning assay. The preference between the two odorants did not differ significantly from zero in either species (as intended with our choice of odour concentrations), and, critically, no significant difference was detected between species (Fig. 2c).

Together, this established conditions for the learning experiments equilibrating the two species with respect to motivational state, sugar- and odour-preference. We therefore next compared odour-reward associative learning performance between the species under these conditions.

Upon odour-reward training, both species exhibited positive learning indices (LI), demonstrating that they were capable of forming odour–reward associative memories (Fig. 3). Critically, LI values of *D*. *elegans* were significantly lower than for *D*. *melanogaster*, both in the pooled analysis (Fig. 3) and when males and females were analysed separately (Fig. S2a).

**Fig. 3.**
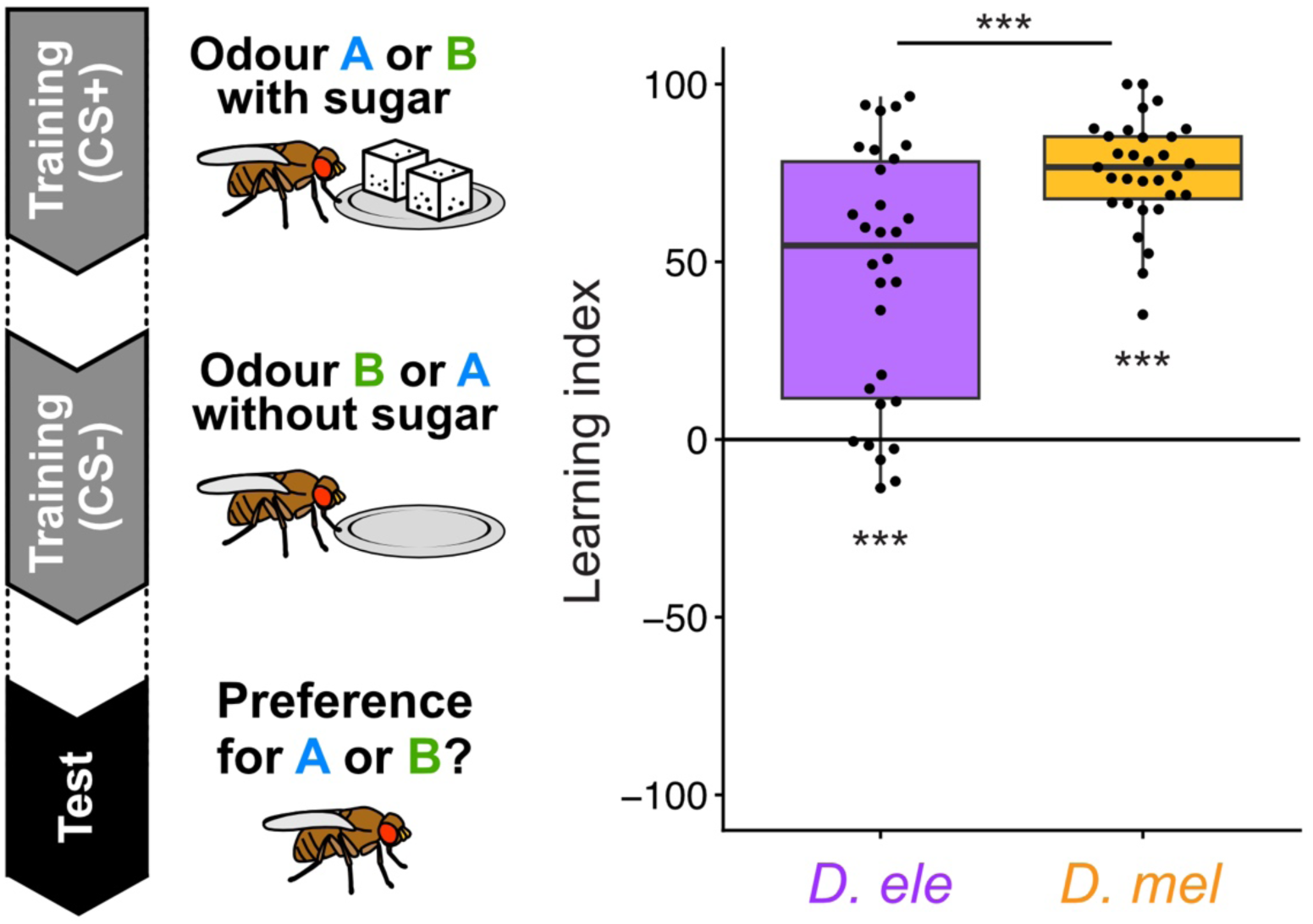
*D*. *elegans* exhibits lower odour-reward associative performance than *D*. *melanogaster* (Wilcoxon rank-sum test, p = 2.01 × 10⁻⁴; asterisks placed above bars). The experimental design of the assay is shown to the left. LI values were significantly greater than zero in both species (Wilcoxon signed-rank test: *D*. *elegans*, p = 2.07 × 10⁻⁵, N = 30; *D*. *melanogaster*, p = 2.46 × 10⁻⁶, N = 31; asterisks placed below individual bars). Asterisks denote statistically significant differences (*p < 0.05, **p < 0.01, ***p < 0.001) and n.s. denotes non-significance.

We next compared colour-reward associative learning abilities between both species, using blue and green ambient light as conditioned stimuli. To exclude effects of brightness, light intensities were adjusted separately for each species such that blue and green stimuli had equivalent subjective brightness (see Material and Methods, Fig. S1). In both species, the preference between blue *versus* green light were significantly greater than zero, indicating a moderate preference for blue in both species, but critically, no significant interspecific difference was detected (Fig. 4a). We therefore compared colour-reward associative learning performance between the species using these visual stimuli. The LIs of *D*. *elegans* were significantly above zero, demonstrating its ability to form colour–reward associative memories (Fig. 4b). In contrast, the LIs of *D*. *melanogaster* did not differ significantly from zero, providing no evidence of colour learning under the present experimental conditions. Critically, the direct comparison of LI values revealed that *D*. *elegans* exhibited significantly higher learning performance than *D*. *melanogaster* in the pooled analysis. When males and females were analysed separately, a significant interspecific difference was detected in males, whereas females showed a similar trend, although the difference was not statistically significant (Fig. S3a). Interestingly, a species-specificity was observed only when the blue light was the rewarded stimulus, but not when the green light was rewarded (Fig. S3b), suggesting that it may be the blue-reward association ability, rather than the green-reward association ability, that differs between both species.

**Fig. 4.**
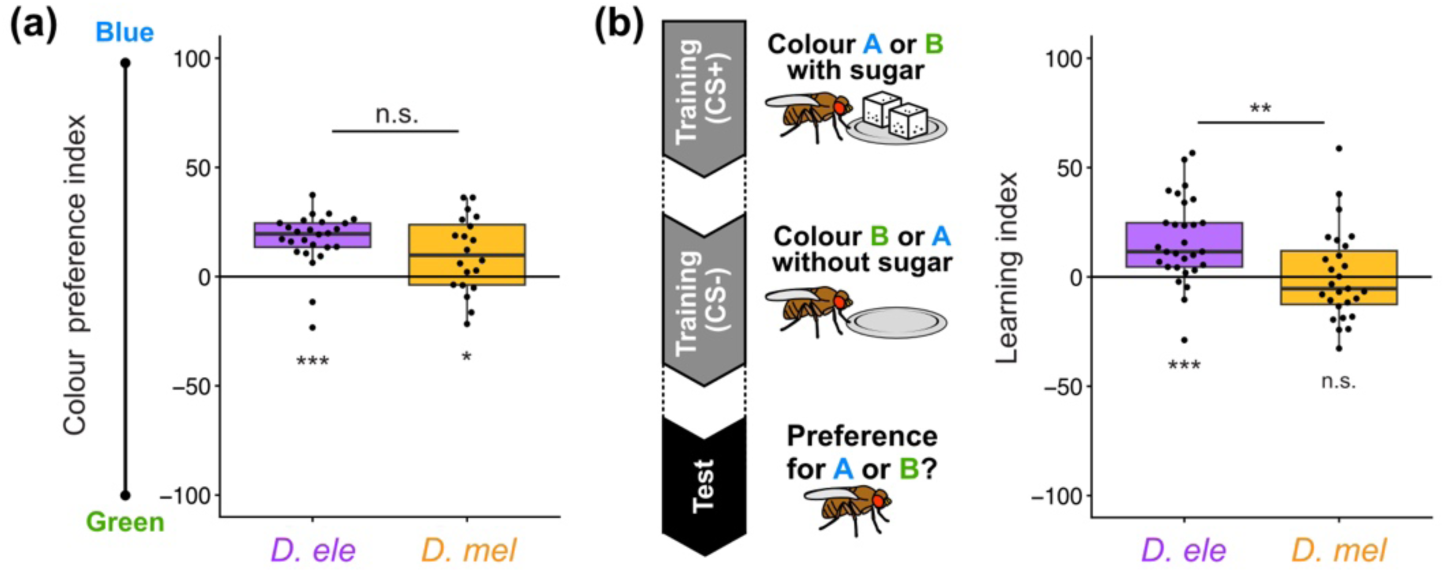
(a) Innate coloured-light preference between blue *versus* green light. Both species exhibited a significant preference for blue light (Wilcoxon signed-rank test: *D*. *elegans*, p = 2.26 × 10⁻⁴, N = 26; *D*. *melanogaster*, p = 0.0152, N = 20; asterisks placed below individual bars). No significant difference was detected between species (Wilcoxon rank-sum test, p = 0.1661; n.s. placed above bars). (b) *D*. *elegans* exhibits higher colour-reward associative performance compared with *D*. *melanogaster* (Student’s t-test, p = 0.003496; asterisks placed above bars). The experimental design of the assay is shown to the left. The LIs of *D*. *elegans* were significantly greater than zero, whereas the LIs of *D*. *melanogaster* did not differ from zero (Wilcoxon signed-rank test: *D*. *elegans*, p = 3.42 × 10⁻⁶, N = 29; *D*. *melanogaster*, p = 0.951, N = 27; asterisks and n.s. placed below individual bars).

## 4. Discussion

We compared associative learning abilities between the flower-feeding species *D. elegans* and the fruit-feeding species *D. melanogaster*. We found that *D. elegans* exhibited lower learning performance for odours but higher learning performance for colours than *D. melanogaster.* Thus, in otherwise similar tasks, under conditions of equilibrated motivation, odour- and colour-choice, and using the same sugar reward, both species differ in the eligibility of odour and colour cues for the establishment of predictive memories. This provides a study case of species-specificity in what is known in experimental psychology as the ‘Garcia effect’: Garcia and Koelling (1966) showed that an audiovisual stimulus is easily associated with an electric shock but not with sickness, while a gustatory stimulus, conversely, is easily associated with sickness but not with electric shock ^24^. Such ‘preparedness’ challenged the idea of generality of associability. Our results extend this notion as they suggest that such preparedness is itself species-specific, and furthermore suggest that biological ‘intelligence’ is not general.

One explanation for why the associative eligibility of colour and odour is doubly dissociated between *D. elegans* and *D. melanogaster* is that the ecological importance of these modalities differs between them. *D. melanogaster* primarily exploits fermenting fruits and is known to locate such resources using volatile compounds together with achromatic visual cues ^32,33^. In contrast, *D. elegans* is strongly reliant on flowers that exhibit diverse visual characteristics ^27^. Indeed, visual signals of flowers have been shown in many plant species to correlate with floral age, pollination status, and reward availability ^15,41^. Flower-visiting animals can therefore use such information to improve foraging efficiency ^11,14^. Under such ecological conditions, selection may favour reliance on colour information and colour-based associative learning. Our results thus encourage further analysis of the sensory cues used by *D. elegans* during natural host and resource choice.

The relatively poor odour-learning performance of *D. elegans* may in turn indicate that odour information is less important in its ecology than is the case for *D. melanogaster*. Interestingly, several olfactory receptor genes have been lost in both *D. elegans* and its flower-feeding sister species *D. gunungcola* ^42^ (Fig. 1), broadly consistent with reduced reliance on olfactory information in the *elegans* group.

At the circuit level it is established for *D. melanogaster* that odour-reward and colour-reward association formation takes place in the mushroom bodies, evolutionarily highly conserved central-brain structures of arthropods ^25,26,43^. In *D. melanogaster*, the mushroom bodies receive input from about the same number of olfactory and visual projection neurons; however, the number of synapses from these projection neurons towards the mushroom bodieś intrinsic neurons, and the number of mushroom body intrinsic neurons handling olfactory and visual signals, is about 10 times higher for olfaction than for vision ^44^. We speculate that this relative ‘numerical disinterest’ in visual processing in *D. melanogaster* might be less pronounced in *D. elegans*, allowing for relatively better colour learning in this flower-feeding species.

The present results should be interpreted with at least two caveats. First, we examined only a single form of associative learning, namely classical conditioning with a sugar reward. Whether similar patterns occur for example for other rewards or for longer-term memories, as well as in aversive, operant or non-associative learning paradigms remains unknown. Second, our comparison was restricted to two species, and broader comparative analyses across multiple flower-feeding and non-flower-feeding *Drosophila* species will be necessary to ascertain the generality of the present findings.

## Data accessibility

All data and R code generated or analysed during this study are available on Zenodo at https://doi.org/10.5281/zenodo.22807867.

## Acknowledgements

We acknowledge Anna Ciuraszkiewicz, Bettina Kracht, Melissa Prothmann, Diana Walther, Canan Nazarov and Isabel Walther for expert technical assistance in *Drosophila* husbandry and stock maintenance, and Ulrich Thomas for kindly providing fly food. We also thank Haruka Yamazaki and Munehiro Katsura for valuable discussions. We thank the Ehime-Fly stock centre for providing *Drosophila* stocks.

## Authors’ contributions

K. K.: Data curation, Formal analysis, Investigation, Funding acquisition, Visualization, Writing – original draft.

C. K.: Investigation, Data curation, Formal analysis, Methodology, Supervision.

T. T.: Conceptualization, Supervision, Writing – review & editing.

B. G.: Conceptualization, Supervision, Writing – review & editing.

Y. I.: Conceptualization, Formal analysis, Funding acquisition, Project administration, Supervision, Visualization, Writing – original draft, Writing – review & editing.

## References

1. Jones PL, Agrawal AA. Learning in insect pollinators and herbivores. Annu Rev Entomol. 2017;62(1):53–71. doi:10.1146/annurev-ento-031616-034903

2. Kamil AC, Roitblat HL. The ecology of foraging behavior: implications for animal learning and memory. Annu Rev Psychol. 1985;36(1):141–169. doi:10.1146/annurev.ps.36.020185.001041

3. Mery F. Natural variation in learning and memory. Current Opinion in Neurobiology. 2013;23(1):52–56. doi:10.1016/j.conb.2012.09.001

4. Ratcliffe JM, Fenton MB, Galef BG. An exception to the rule: common vampire bats do not learn taste aversions. Animal Behaviour. 2003;65(2):385–389. doi:10.1006/anbe.2003.2059

5. Liu Y, Jones CD, Day LB, Summers K, Burmeister SS. Cognitive phenotype and differential gene expression in a hippocampal homologue in two species of frog. Integr Comp Biol. 2020;60(4):1007–1023. doi:10.1093/icb/icaa032

6. Krenn HW, Plant JD, Szucsich NU. Mouthparts of flower-visiting insects. Arthropod Structure & Development. 2005;34(1):1–40. doi:10.1016/j.asd.2004.10.002

7. Wäckers FL, Romeis J, Van Rijn P. Nectar and pollen feeding by insect herbivores and implications for multitrophic interactions. Annu Rev Entomol. 2007;52(1):301–323. doi:10.1146/annurev.ento.52.110405.091352

8. Xiao L, Labandeira C, Dilcher D, Ren D. Florivory of early Cretaceous flowers by functionally diverse insects: implications for early angiosperm pollination. Proc R Soc B. 2021;288(1953):20210320. doi:10.1098/rspb.2021.0320

9. Ollerton J. Pollinator diversity: distribution, ecological function, and conservation. Annu Rev Ecol Evol Syst. 2017;48(1):353–376. doi:10.1146/annurev-ecolsys-110316-022919

10. Potts SG, Biesmeijer JC, Kremen C, Neumann P, Schweiger O, Kunin WE. Global pollinator declines: trends, impacts and drivers. Trends in Ecology & Evolution. 2010;25(6):345–353. doi:10.1016/j.tree.2010.01.007

11. Lunau K, Maier EJ. Innate colour preferences of flower visitors. J Comp Physiol A. 1995;177(1):1–19. doi:10.1007/BF00243394

12. Nicolson SW. Sweet solutions: nectar chemistry and quality. Phil Trans R Soc B. 2022;377(1853):20210163. doi:10.1098/rstb.2021.0163

13. Raguso RA. Wake up and smell the roses: The ecology and evolution of floral scent. Annual Review of Ecology, Evolution, and Systematics. 2008;39:549–569.

14. van der Kooi CJ, Dyer AG, Kevan PG, Lunau K. Functional significance of the optical properties of flowers for visual signalling. Annals of Botany. 2019;123(2):263–276. doi:10.1093/aob/mcy119

15. van der Kooi CJ, Reuvers L, Spaethe J. Honesty, reliability, and information content of floral signals. iScience. 2023;26(7):107093. doi:10.1016/j.isci.2023.107093

16. Guo M, Du L, Chen Q, et al. Odorant receptors for detecting flowering plant cues are functionally conserved across moths and butterflies. Molecular Biology and Evolution. 2021;38(4):1413–1427. doi:10.1093/molbev/msaa300

17. Sharkey CR, Powell GS, Bybee SM. Opsin evolution in flower-visiting beetles. Front Ecol Evol. 2021;9. doi:10.3389/fevo.2021.676369

18. Valencia-Montoya WA, Liénard MA, Rosser N, et al. Infrared radiation is an ancient pollination signal. Science. 2025;390(6778):1164–1170. doi:10.1126/science.adz1728

19. Menzel R. The honeybee as a model for understanding the basis of cognition. Nat Rev Neurosci. 2012;13(11):758–768. doi:10.1038/nrn3357

20. Wright GA, Schiestl FP. The evolution of floral scent: the influence of olfactory learning by insect pollinators on the honest signalling of floral rewards. Functional Ecology. 2009;23(5):841–851. doi:10.1111/j.1365-2435.2009.01627.x

21. Riffell JA, Alarcón R, Abrell L, Davidowitz G, Bronstein JL, Hildebrand JG. Behavioral consequences of innate preferences and olfactory learning in hawkmoth– flower interactions. Proceedings of the National Academy of Sciences. 2008;105(9):3404–3409. doi:10.1073/pnas.0709811105

22. Robert T, Tarapata K, Nityananda V. Learning modifies attention during bumblebee visual search. Behav Ecol Sociobiol. 2024;78(2):22. doi:10.1007/s00265-024-03432-z

23. Greggers U, Menzel R. Memory dynamics and foraging strategies of honeybees. Behav Ecol Sociobiol. 1993;32(1):17–29. doi:10.1007/BF00172219

24. Garcia J, Koelling RA. Relation of cue to consequence in avoidance learning. Psychonomic Science. 1966;4(1):123–124. doi:10.3758/BF03342209

25. Modi MN, Shuai Y, Turner GC. The *Drosophila* mushroom mody: From architecture to algorithm in a learning circuit. Annu Rev Neurosci. 2020;43(1):465–484. doi:10.1146/annurev-neuro-080317-0621333

26. Vogt K, Schnaitmann C, Dylla KV, et al. Shared mushroom body circuits underlie visual and olfactory memories in *Drosophila*. eLife. 2014;3:e02395. doi:10.7554/eLife.02395

27. Ishikawa Y, Kimura MT, Toda MJ. Biology and ecology of the oriental flower-breeding *Drosophila elegans* and related species. Fly (Austin). 2022;16(1):207–220. doi:10.1080/19336934.2022.2066953

28. Prud’homme B, Gompel N, Rokas A, et al. Repeated morphological evolution through cis-regulatory changes in a pleiotropic gene. Nature. 2006;440(7087):1050–1053. doi:10.1038/nature04597

29. Kimura M, Hirai Y. Daily activity and territoriality of *Drosophila elegans* in Sukarami, West Sumatra, Indonesia. Tropics. 2001;10(3):489–495. doi:10.3759/tropics.10.489

30. Suwito A, Ishida T, Hattori K, Kimura M. Territorial and mating behaviours of two flower-breeding *Drosophila* species, *D. elegans* and *D. gunungcola* (Diptera: Drosophilidae) at Cibodas, West Java, Indonesia. Treubia. 2012;39:77–85. doi:10.14203/treubia.v39i0.24

31. Hirai Y, Kimura MT. Incipient reproductive isolation between two morphs of *Drosophila elegans* (Diptera: Drosophilidae). Biological Journal of the Linnean Society. 1997;61(4):501–513. doi:10.1006/bijl.1996.0133

32. Frye MA, Tarsitano M, Dickinson MH. Odor localization requires visual feedback during free flight in *Drosophila melanogaster*. J Exp Biol. 2003;206(Pt 5):843–855. doi:10.1242/jeb.00175

33. Lebreton S, Becher PG, Hansson BS, Witzgall P. Attraction of *Drosophila melanogaster* males to food-related and fly odours. Journal of Insect Physiology. 2012;58(1):125–129. doi:10.1016/j.jinsphys.2011.10.009

34. Thiagarajan D, Eberl F, Veit D, Hansson BS, Knaden M, Sachse S. Aversive bimodal associations differently impact visual and olfactory memory performance in *Drosophila*. iScience. 2022;25(12):105485. doi:10.1016/j.isci.2022.105485

35. Yoshida T, Chen H, Toda MJ, Kimura MT, Davis AJ. New host plants and host plant use for *Drosophila elegans* Bock and Wheeler. Drosoph Inf Serv. 2000;83:18–21.

36. Tully T, Quinn WG. Classical conditioning and retention in normal and mutant *Drosophila melanogaster*. J Comp Physiol. 1985;157(2):263–277. doi:10.1007/BF01350033

37. Colomb J, Kaiser L, Chabaud MA, Preat T. Parametric and genetic analysis of *Drosophila* appetitive long-term memory and sugar motivation. Genes Brain Behav. 2009;8(4):407–415. doi:10.1111/j.1601-183X.2009.00482.x

38. Tempel BL, Bonini N, Dawson DR, Quinn WG. Reward learning in normal and mutant Drosophila. Proceedings of the National Academy of Sciences. 1983;80(5):1482–1486. doi:10.1073/pnas.80.5.1482

39. Scheiner R, Steinbach A, Claßen G, Strudthoff N, Scholz H. Octopamine indirectly affects proboscis extension response habituation in *Drosophila melanogaster* by controlling sucrose responsiveness. Journal of Insect Physiology. 2014;69:107–117. doi:10.1016/j.jinsphys.2014.03.011

40. Uchizono S, Tanimura T. Genetic variation in taste sensitivity to sugars in *Drosophila melanogaster*. Chem Senses. 2017;42(4):287–294. doi:10.1093/chemse/bjw165

41. Ruxton GD, Schaefer HM. Floral colour change as a potential signal to pollinators. Current Opinion in Plant Biology. 2016;32:96–100. doi:10.1016/j.pbi.2016.06.021

42. Negi A, Liao BY, Yeh SD. Long-read-based genome assembly of *Drosophila gunungcola* reveals fewer chemosensory genes in flower-breeding species. Genome Biol Evol. 2023;15(3):evad048. doi:10.1093/gbe/evad048

43. Strausfeld NJ, Wolff GH, Sayre ME. Mushroom body evolution demonstrates homology and divergence across Pancrustacea. eLife. 2020;9:e52411. doi:10.7554/eLife.52411

44. Li F, Lindsey JW, Marin EC, et al. The connectome of the adult *Drosophila* mushroom body provides insights into function. eLife. 2020;9:e62576. doi:10.7554/eLife.62576

