## Supplementary Material for "Distinct associative learning abilities for colour and odour in the flower-feeding *Drosophila elegans* and the fruit-feeding *Drosophila melanogaster*"

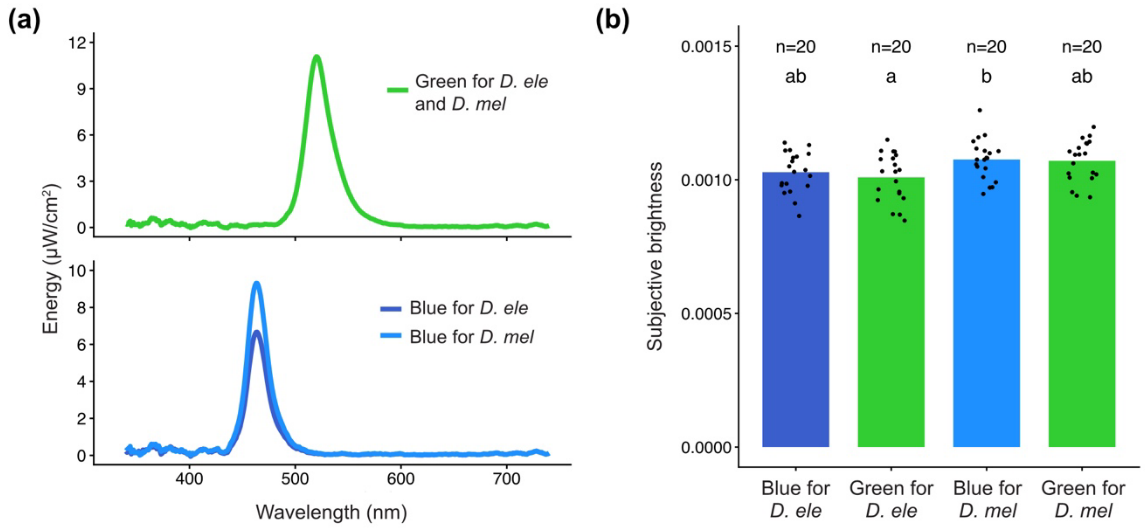


Fig. S1. (a) The emission spectra of the LED light sources used in this study. Curves represent the mean emission intensity at each wavelength based on five repeated measurements of each device. (b) The subjective brightness of the LED light sources used in this study. Dots represent five replicate measurements, bars indicate the mean. Different letters above bars indicate statistically significant differences (see Table S1).

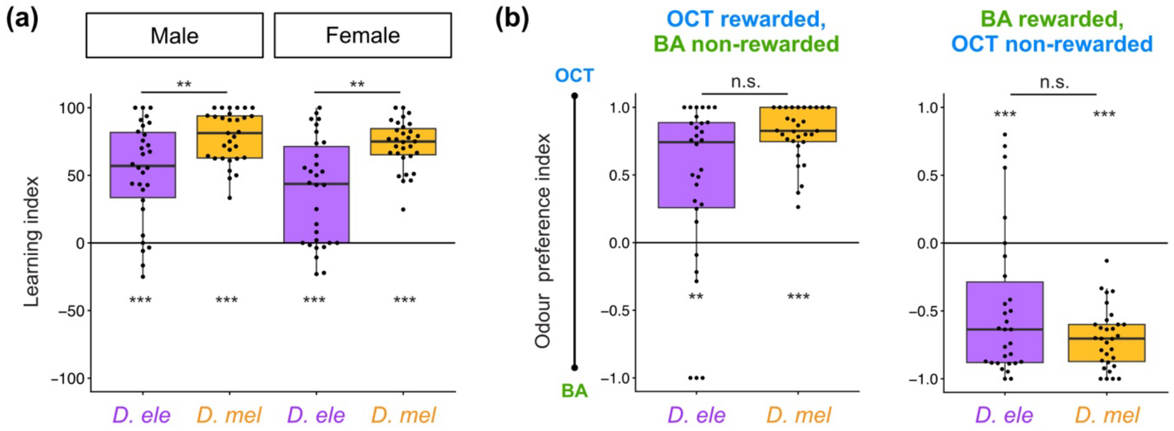


Fig. S2. (a) Learning indices (LIs) from Fig. 3, separated by sex. In both sexes, *D. melanogaster* exhibited significantly higher LI values than *D. elegans* (Wilcoxon rank-sum test with Holm correction: females, p = 0.0011; males, p = 0.00442). Both male and female flies of both species exhibited significantly positive LI values (Wilcoxon signed-rank test with Holm correction: *D. elegans* males, $p = 2.16 \times 10^{-5}$; *D. melanogaster* males, $p = 4.83 \times 10^{-6}$; *D. elegans* females, $p = 1.54 \times 10^{-4}$; *D. melanogaster* females, $p = 4.83 \times 10^{-6}$). (b) Odour preference indices for octanol (OCT) *versus* benzaldehyde (BA) underlying the learning indices in Fig. 3. No significant interspecific difference was detected (Wilcoxon rank-sum test with Holm correction: OCT rewarded, p = 0.0528; BA rewarded, p = 0.302). Both species exhibited a significant preference for the sucrose-paired odour regardless of the chemical identity of the rewarded odour (Wilcoxon signed-rank test with Holm correction: *D. elegans*, OCT rewarded, p = 0.00296; *D. melanogaster*, OCT rewarded, $p = 4.47 \times 10^{-6}$; *D. elegans*, BA rewarded, $p = 7.80 \times 10^{-4}$; *D. melanogaster*, BA rewarded, $p = 4.47 \times 10^{-6}$).

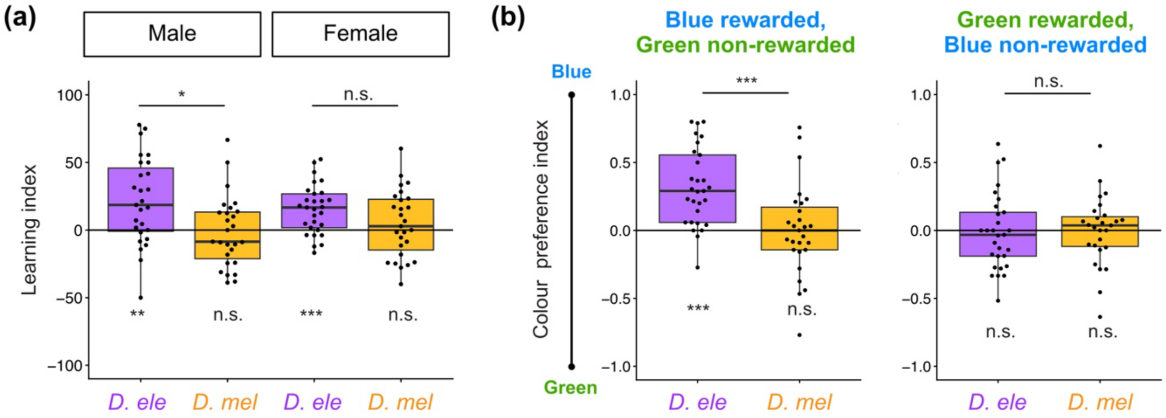


Fig. S3. (a) Learning indices (LIs) from Fig. 4b, separated by sex. In males, *D. elegans* exhibited significantly higher LIs than *D. melanogaster*, whereas no significant difference between species was detected in females (Welch's t-test with Holm correction: males, $p = 0.0116$; females, $p = 0.0726$). Both male and female of *D. elegans* exhibited significantly positive LI values, whereas neither male nor female of *D. melanogaster* showed a significant deviation from zero (one-sample t-test with Holm correction: *D. elegans* males, $p = 0.00522$; *D. melanogaster* males, $p = 0.771$; *D. elegans* females, $p = 3.91 \times 10^{-4}$; *D. melanogaster* females, $p = 0.683$). (b) Colour preference indices for blue *versus* green underlying the learning indices in Fig. 4b. *D. elegans* exhibited a significantly higher preference index than *D. melanogaster* when blue light was paired with reward, whereas no significant interspecific difference was detected when green light was paired with reward (Welch's t-test with Holm correction: blue rewarded, $p = 7.76 \times 10^{-4}$; green rewarded, $p = 0.896$). *D. elegans* exhibited a significant preference for the reward-paired colour when blue was rewarded, whereas *D. melanogaster* showed no significant preference in either training condition (one-sample t-test with Holm correction: *D. elegans*, blue rewarded, $p = 6.85 \times 10^{-6}$; *D. melanogaster*, blue rewarded, $p = 1.0$; *D. elegans*, green rewarded, $p = 1.0$; *D. melanogaster*, green rewarded, $p = 1.0$).

| Variable | F (df1, df2) | ANOVA p | Tukey comparison | Difference | Adjusted p |
|---|---|---|---|---|---|
| Colour*Species | 3.293(3,76) | 0.025 | Blue for *D. mel* – Blue for *D. ele* | 4.737395E-05 | 0.252 |
| | | | Green for D. ele – Blue for *D. ele* | -1.938640E-05 | 0.871 |
| | | | Green for *D. mel* – Blue for *D. ele* | 4.262280E-05 | 0.343 |
| | | | Green for *D. ele* – Blue for *D. mel* | -6.676035E-05 | 0.0499988 |
| | | | Green for *D. mel* – Blue for *D. mel* | -4.751150E-06 | 0.998 |
| | | | Green for *D. mel* – Green for *D. ele* | 6.200920E-05 | 0.0782 |

Table S1. Statistical analysis of subjective brightness data shown in Fig. S1 (one-way ANOVA followed by Tukey's post hoc test).

| | 0 h starvation | | 24 h starvation | | 48 h starvation |
|---|---|---|---|---|---|
| | *D. ele* | *D. mel* | *D. ele* | *D. mel* | *D. ele* |
| p-value | 0.0914 | 0.564 | $1.72 \times 10^{-5}$ | $1.30 \times 10^{-4}$ | 0.00218 |

Table S2. Statistical analysis of sucrose preference indices from Fig. 2b, reporting tests against zero for each species and starvation condition (Wilcoxon signed-rank tests with Holm correction for multiple comparisons).

| | | 0 h starvation | | 24 h starvation | | 48 h starvation |
|---|---|---|---|---|---|---|
| | | *D. ele* | *D. mel* | *D. ele* | *D. mel* | *D. ele* |
| 0 h starvation | *D. ele* | | | | | |
| | *D. mel* | 1.0 | | | | |
| 24 h starvation | *D. ele* | 0.0394 | 0.0153 | | | |
| | *D. mel* | 0.0091 | 0.0036 | 1.0 | | |
| 48 h starvation | *D. ele* | 0.0070 | 0.0025 | 1.0 | 1.0 | |

Table S3. Pairwise comparisons of sucrose preference indices from Fig. 2b, reporting tests among species × starvation-duration groups (an aligned rank transform (ART) ANOVA adjusted using the Holm method).
